# Random burst sensing of neurotransmitters

**DOI:** 10.1101/607077

**Authors:** P. Read Montague, Terry Lohrenz, Jason White, Rosalyn J Moran, Kenneth T. Kishida

## Abstract

We introduce a random sensing approach to neurotransmitter detection that provides concurrent, co-localized detection of dopamine, serotonin, norepinephrine and pH. The approach generates high quality out-of-sample predictions at 10 milliseconds per estimate. Similar high-quality estimates result when the data are down-sampled suggesting that even more dramatic speedups are possible. The method also works using electrophysiological probes in routine use in clinical preparations thus transforming these and similar electrodes into ultra-fast sources of multi-transmitter information.

---

We introduce a new approach for co-localized, sub-second detection of dopamine, serotonin, and norepinephrine. The method derives from a recent innovation called elastic net electrochemistry^1,2,3^, which has been used to make the first sub-second recordings of dopamine and serotonin from human brain^1,2^. The method introduced here, which we term random burst sensing (RBS), relies on the application of a randomized, repeating pattern of voltages, and uses the electrode current responses to infer all three neuromodulators concurrently. We further demonstrate that the method works well on standard platinum-iridium electrodes in routine use in human depth recordings, deep brain stimulators, and other commonly used electrodes^4^ – thus augmenting the functional repertoire of normal electrodes to include fast neuromodulator detection.

In an effort to use voltammetry in conscious human brains, we redefined the approach to voltammetry by building a supervised inference approach to estimate neuromodulator concentrations^1,2,3^. The goal was to extract a model *in vitro* that could be trained to predict and generalize to novel neurotransmitter measurements and novel (but similar) electrodes *in vivo*. We called this approach Elastic net electrochemistry and showed that it works at sub-second rates, displays robustness against pH changes (see fig. 2 Kishida et al., 2016) and estimates serotonin and dopamine concurrently off the same electrode^2^. The same approach can estimate norepinephrine and distinguish it from both dopamine and serotonin^3^. In all that work, the voltage forcing function was a standard triangular waveform 10 milliseconds in duration either followed by a 90 millisecond ‘wait’ period^5^ (100 msec duty cycle^1,2^) or repeating contiguously^3^.

However, two features of elastic net electrochemistry suggested that a useful, but radical departure from this approach was possible: (1) significant random down-sampling of the transformed current time series produced excellent multi-transmitter prediction models^3^, and (2) the concentration-dependent encoding responses of the electrode were ***not solely concentrated at the oxidation potential*** of the neuromodulators but spread coherently, but ‘wiggling’ throughout the entire current time series^2,3^. These observations suggested the presence of redundant, stable concentration information spread throughout the current time series sufficient to extract excellent concentration-prediction models for multiple, coactive neuromodulators, and at potentially faster timescales.

Figure 1a shows the workflow for random burst sensing. Voltages throughout the range containing the oxidation potential of dopamine and serotonin were quantized into 40 segments and randomized. This pattern was repeatedly delivered every 10.3 milliseconds (red inset shows detailed structure of random burst in this example). The workflow bifurcates where the top sequences uses all data in each 10.3 millisecond burst and the bottom sequence uses only a random 10% of the data. Otherwise the steps are identical as indicated. The final step for both branches is to extract a constrained regression model using either the elastic net^6^ or LASSO regression^7^. This is all carried out in a flow cell where the concentration of analytes can be controlled and we show results for a carbon fiber electrode compared to our usual stainless steel ground electrode as reported previously^1,2^. Figure 1b,c shows specifically the out-of-sample predictions for the cross-validated models so extracted for dopamine and serotonin. The performance of each class of model (i.e. using 100% or 10% of the data) is shown in two ways: (1) a plot summarizing average out-of-sample predictions across a range, and (2) a plot showing the model output to step changes in either dopamine or serotonin.

**Figure 1.**
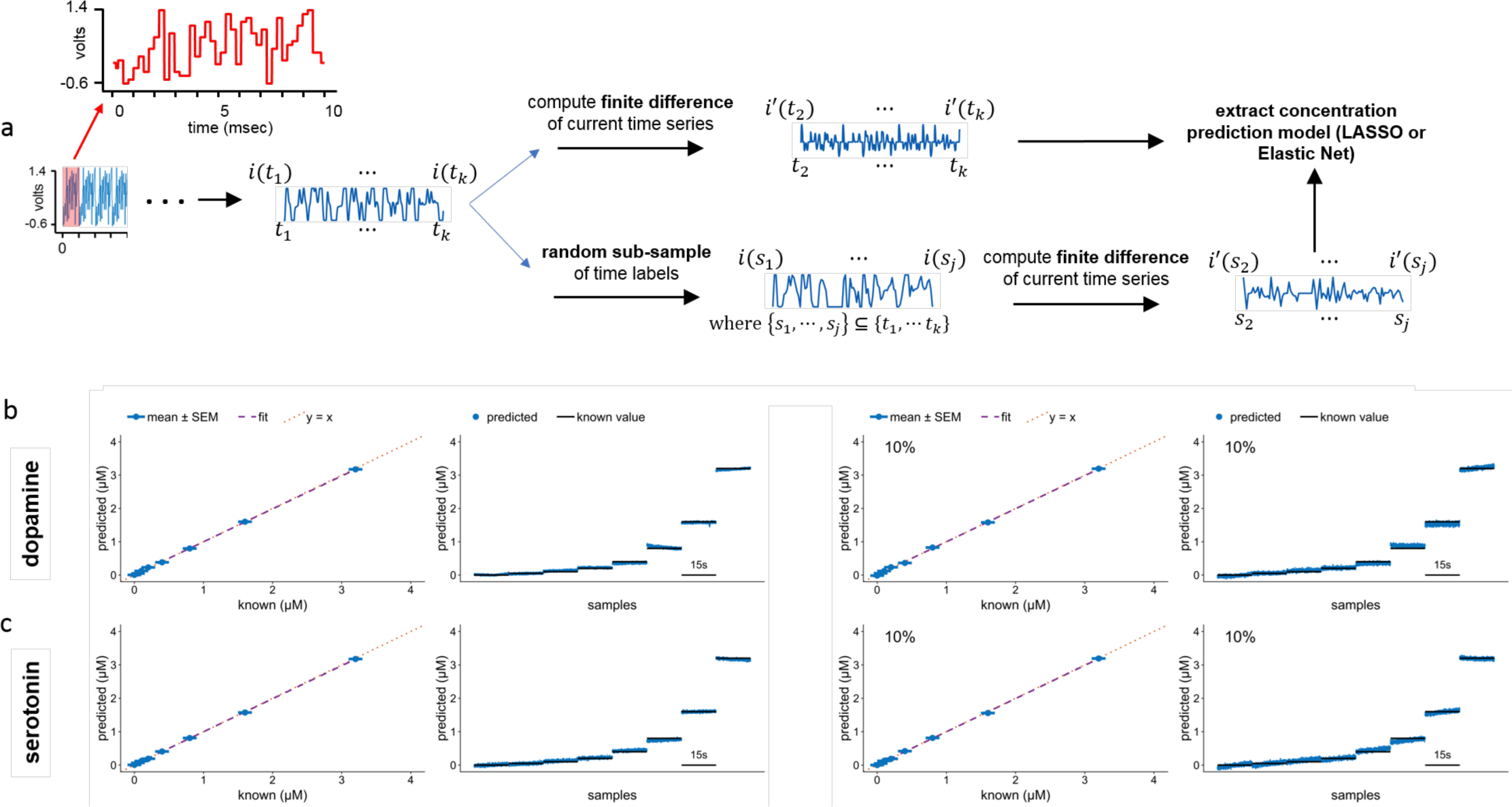
**a)** Workflow for extracting neuromodulator prediction models from randomized voltage waveform (red trace, inset). Workflow bifurcates for using 100% of the measured data versus a dramatically downsampled 10%. Note that after downsampling the finite difference trace is still computed before entering the data into either elastic net or LASSO regression. **b)** Dopamine predictions using random burst sensing. Average out-of-sample predictions shown next to model predictions to stepwise changes in dopamine. **c)** Serotonin predictions using random burst sensing. 100% and 10% conditions labeled. Note that 10% of measured data represents only 1 millisecond of data. Although not shown, even less data than 1 millisecond can be used to produce excellent out-of-sample models.

We then sought to determine whether the random burst method could disambiguate four critical neuromodulatory chemicals – using concentration mixtures comprising one of the 4 analytes. Figure 2a shows the results of a random burst-derived model in the presence of 4 analytes – dopamine, serotonin, norepinephrine, and 5-hydroxyindoleacetic-acid (5HIAA). The separation of dopamine and norepinephrine by voltammetric methods was previously not thought possible, but we suspect that belief may have emerged from a focus solely on the oxidation potential for these two neurotransmitters, which share nearly identical waveforms near those potentials. The top panel shows the 10.3 millisecond predictions for each transmitter as a colored dot, the horizontal black lines show actual level, and the bottom panel shows the average predictions for each 15 second bin. To ensure that temporally co-localized fluctuations in neuromodulatory concentrations could be disentangled we tested the model using mixtures of 5HT and 5-HIAA (Fig 2b). Here again we showed that despite very large potential contamination from 5-HIAA, the underlying, true levels of serotonin could be selectively identified.

**Figure 2.**
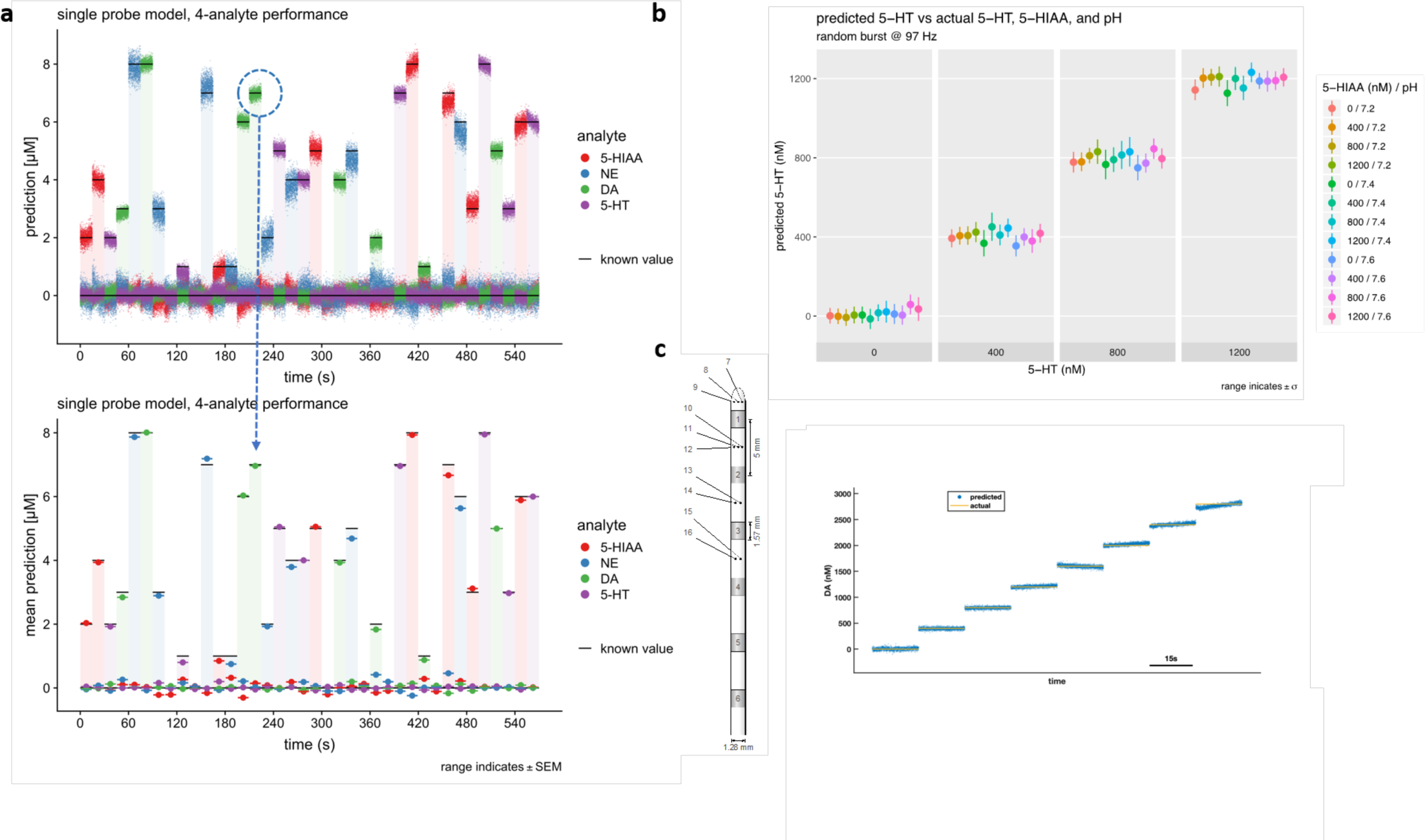
**a)** Multi-transmitter model using 10.3 msec estimates. Top panel: Predictions for 4 analytes at 10.3 msec per estimate. Black lines are actual level. **Bottom panel**: Averages for each 15 second window. Black lines are actual level. **b)** 4 serotonin levels versus ranges of 5-hydroxyindoleacetic acid and pH. Each prediction is for a 10.3 msec model. **c)** Random burst electrochemistry on commercial electrode made of platinum iridium using 10.3 msec random burst activation. Here we show proof-of-principle result for step changes in dopamine levels (out-of-sample). Similarly good models were extracted for the other 3 analytes in fig 2a (not shown).

Finally, we sought to determine whether the method could be applied to electrodes used in routine clinical practice. We trained and tested the random burst-derived model using flow-cell current recordings on a platinum iridium electrode. We show excellent out-of-sample predictions for dopamine on this depth electrode (fig. 2c). Random burst sensing was also successfully applied to 4 analytes as in figure 2a (not shown). This proof-of-principle demonstration on a commercially available electrode in common use suggests a potentially transformative use of this new approach.

## Acknowledgements

This work was supported by a Principal Research Fellowship from The Wellcome Trust (PRM), Virginia Tech (PRM, KTK); Wake Forest School of Medicine (KTK); and the National Institutes of Health: R01 NS092701 (PRM, KTK), KL2 5KL2TR00142, (KTK), UL1TR001420 (KTK), and P50AA026117 (KTK).

## Methods

### Equipment

We perform random burst sensing (RBS) of neurotransmitters *in vitro* using both commercially-available platinum-iridium macro-micro depth electrodes (MM16A-SP05X-000; AdTech) and extended carbon-fiber microsensors constructed in-house using the same materials and procedures as those we use *in vivo* ^1,2^. When collecting data *in vitro* from the commercial electrode, we select one microwire contact (Pt/Ir, 0.051 mm diameter) as the working electrode, and a separate microwire contact ∼5 mm away as the reference. By contrast, the carbon-fiber microsensor utilizes a stainless-steel reference located ∼10 mm from the working electrode. The non-working end of the probe is held in place by a guide tube and the working end protrudes into a small flow cell. For the larger commercial probe, we use the tip of a plastic transfer pipette (13-711-9AM; Fischer Scientific) for both the guide tube and the flow cell. For the carbon-fiber probe, we use a microelectrode insertion tube (FC1036; FHC) as the guide tube and a glass capillary tube (PG52165-4; World Precision Instruments) as the flow cell. The commercial probe utilizes a pigtail adaptor (macro: L-SRL-6DIN, micro: L-SRL-10DIN; AdTech) to connect the various macro- and micro-contacts. The carbon-fiber probe is constructed with standard gold-plated pin connectors for working and reference electrodes.

Due to differences in impedance and electrode composition, we do not mix data acquired from the commercial probes with those acquired from the carbon-fiber probes; rather, we build separate models for each probe type (see Fig. 2 main text). Unless otherwise stated, the following details are identical regardless of probe selection. The working and reference electrodes connect to an electrochemistry-capable headstage (CV-7B/EC; Axon Instruments) modified by the manufacturer to accommodate current responses of ±2 µA when operating in voltage clamp mode. The connection is made using a commercial three-conductor shielded microelectrode cable, the same model used to connect monitoring and electrochemical electrodes during DBS implantation surgery (FC1020; FHC). Working and reference leads are connected to the probe, and shield ground is connected to the guide cannula with an alligator clip. We built an adaptor that connects the headstage working and reference ports to a female receptacle compatible with the cable (5-pin DIN, 240º). We connect the shielding ground the ground connection on the rear of the amplifier.

The headstage connects to a signal amplifier / headstage controller (Multiclamp 700B; Axon Instruments) which connects to a digital acquisition (DAQ) system (Digidata 1550B; Axon Instruments). Both the amplifier and the DAQ connect to the recording computer (MacBook Pro; Apple) via USB and are controlled by software: the amplifier by Axon MultiClamp 700B Commander version 2.2.2.2 (Axon Instruments), and the DAQ by Axon Clampex version 10.7.0.3 (Axon Instruments). The recording computer uses the Windows 10 Enterprise operating system (Microsoft).

### Random Burst Forcing Function and Acquisition Protocol

To build the RBS forcing function, we started by discretizing the 1000-sample triangle portion of the FSCV waveform we use *in vivo* ^1^ into 40 constant-voltage steps ranging from −0.6 V to 1.4 V, each 25 samples in length. We shuffled the order of the voltage steps and wrote them to a tab-delimited file format (.ATF) which can be interpreted by the Clampex software.

We loaded the file into Clampex and configured it to use a sampling rate of 100,000 Hz, yielding a random burst “command” waveform with 1000 samples of 10-ms duration consisting of 250-µs constant-voltage steps. When configured to repeat the command waveform at the maximum-allowed frequency, Clampex enforces a minimum hold time before and after each repetition. In our case, this equals a total of 32 extra samples (320 extra microseconds) which yields a repetition rate of approximately 96.90 Hz. Clampex will record a maximum of 10,000 repetitions in a single file, or approximately 103.2 seconds per recording (i.e. per concentration – see below). The acquisition protocol was saved to disk for future use.

Using the Multiclamp Commander software, we configured the headstage to operate in voltage-clamp mode, which holds the potential difference between the reference and working ports at the command voltage and sends the resulting current response back to the DAQ from which it can be recorded by Clampex. We set the headstage parameters to allow a command signal of ±2 V and a current response of ±2 µA. To match our *in-vivo* setup, we configured the hardware filter on the current response channel to an 8-pole Bessel low-pass filter with a cut-off frequency of 4 kHz. The headstage configuration was saved to disk for future use.

### Solution Preparation

We prepare solutions of dopamine, serotonin, norepinephrine, and 5-hydroxyindoleacetic acid in phosphate-buffered saline (PBS) by first dissolving powdered reagents into 0.1N hydrochloric acid to form 10 mM stock solutions, aliquots of which are frozen for later use. We then prepare a 50 µM starting solution by a series of dilutions in 1× PBS at pH 6.8 from the aliquots. The solutions used are created by diluting one or more starting solutions with 1× PBS until the desired concentrations are reached. For our initial analyte analysis (dopamine and serotonin) we use concentrations from 0 – 3 µM (Fig. 1). For our mutli-analyte analysis we use concentrations from 0 – 8 µM (Fig. 2). The pH of the solutions is achieved by using NaOH or HCl to adjust the pH of the 1× PBS used in the final dilution. During recording, each solution is introduced by syringe through a small filling tube into the glass capillary flow cell containing the microsensor probe. The volume of the flow cell and filling tube is sufficiently small that the solution can be fully replaced with injections of less than 1 mL.

### Recording Procedure

Before data acquisition, we soak the probe in a 70% isopropanol bath for ∼5 minutes to enhance wetting of the electrode surfaces. We then place the probe through the guide cannula into the flow cell and inject ∼5 mL of ultra-pure water followed by ∼5 mL of 1× PBS (pH 7.4) and verify there are no air bubbles on the probe. We verify the probe is connected and working properly by running a standard triangle FSCV protocol (not recorded) and observing a typical current response.

We begin an RBS recording by first evacuating the flow cell and filling tube using suction. We then in inject ∼1 mL of 1× PBS (pH 7.4), evacuate again, and inject the test solution containing a specified concentration of one or more neurotransmitters at a specified pH. We then initiate the recording process in Clampex. We configured the process to begin with a 25.8-second pre-cycle (10-ms triangle waveform, 400 V/s at 96.9 Hz) immediately followed by the full 103.2-second RBS recording. We repeat this procedure for each test solution. The digital data files are stored in the proprietary Axon Binary Format (.ABF) in a directory on the recording computer.

### Data Conditioning

In order to evaluate the performance of the RBS method, we use Matlab 2018 (MathWorks) to process each recording session, i.e. a contiguously-acquired group of recordings taken from one probe exposed to varying concentrations of neurotransmitters and pH levels. We first convert the .ABF recording files into Matlab-compatible files (.MAT) using the abfload utility. ^8^ Each file contains 10,000 current response vectors for the mixture of neurotransmitter concentrations and pH for that recording, and we label each current response vector with the vector of concentrations and pH. To avoid false noise sources such as fluid motion or external interference, we extract a 15-second “stable” window from the recording by finding the minimum total RMS difference between each current response in the window and the median current response for the window. From that portion, we exclude any outlying current responses using Matlab’s isoutlier function and split the data into disjoint training and testing sets by randomly selecting 125 current responses for training and keeping the remainder for out-of-sample testing. We combine the training and testing sets of each recording into training and testing data matrices *X*_*train*_ and *X*_*test*_, and training and testing label matrices *Y*_*train*_ and *Y*_*test*_.

### Sub-sampling

To evaluate the possibility of reducing acquisition time by taking advantage of the temporal coherence of the current responses, we take a random sub-sample of each current response *x*_*i*_ in the data matrices *X*_*train*_ and *X*_*test*_ (Fig. 1, b and c, right panels). In our case, each *x*_*i*_ is a time series with index *t* ∈ {*t*_1_ … *t*_1000_} where *x*_*i*_(*t*_*k*_) is the current value at time *t*_*k*_ relative to the beginning of the current response. We create a global sub-sampling index *s* ⊂ *t* by taking a random sample of *t* without replacement and sorting the result to enforce monotonicity. We then use *s* as the time index to extract the subsampled data matrices 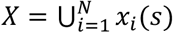. For a given sub-sampling percentage, the sub-sample index *s* is identical for all datasets. We skip this step in the full-sample case, for which *s* = *t*.

### Fitting Elastic Net Regressions

To generate a model that will predict neurotransmitter concentrations 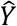 from current responses *X*_*test*_, we first condition the data by computing the finite time difference of the training current responses 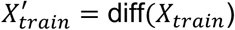, as we find that the regressions fit to 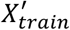 have superior performance to those fit to the undifferentiated data *X*_*train*_. Note that by taking the finite difference we decrease by one the number of samples per observation to 999. We then use the glmnet toolkit for Matlab ^9^ to fit a 10-fold cross-validated multinomial linear regression with LASSO regularization:

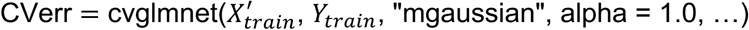

Our invocation of glmnet creates a cross-validated elastic-net regression CVerr having the “multi-response gaussian” objective function ^10,11^:

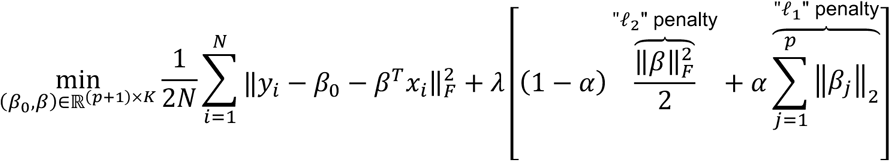

In our case, *N* is the number of observations (125 × number of recordings), *p* is the number of samples per observation (999), and *K* is the number of analytes (neurotransmitters and pH) per label. Each observation 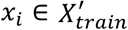 has dimensions 1 × *p*, and each label *y*_*i*_ ∈ *Y*_*train*_ has dimensions 1 × *K*. Here 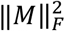 denotes the squared Frobenius norm of an *U* × *V* matrix 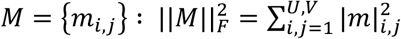. ^12^ Also, the intercept *β*_0_ is a 1 × *K* matrix, the coefficient matrix *β* has dimensions *p* × *K*, and *β*_j_ is a 1 × *K* row in *β*.

The elastic net mixing parameter *α* ∈ [0, 1] controls the relative impacts of the “ℓ_1_” and “ℓ_2_” penalties on *β* indicated above. Selecting *α* = 0 yields a ridge regression in which the *β* coeffects are constrained by their Euclidean magnitude (ℓ_2_) yielding fewer large and more small coefficients. By specifying *α* = 1, we choose a LASSO regression in which the *β* coefficients are constrained by the number of non-zero elements (ℓ_1_) yielding fewer non-zero coefficients.

Glmnet finds the coefficient matrix *β* that satisfies the objective for a range of values for the complexity variable *λ*. To generate predictions for our analyses, we use the “lambda_min” parameter to select the *β* matrix that gives the minimum mean of the squared cross-validated residuals.

The folds used for cross validation are determined randomly by default, though we calculate our own random fold selection in advance. This allows for reproducible comparisons between methods, and a more stable way perform a grid search for the best *α* value should we so desire. Structured approaches to building the folds may yield regressions optimized for different goals, such as concentration interpolation or generalization across probes.

### Prediction Generation and Data Analysis

For the test set, we generate the matrix of predicted concentration and pH values 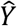 from a cross-validated model using the CVerr object and the finite time difference of the testing current responses 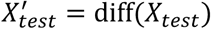 using another glmnet call:

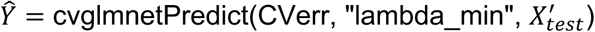

Where 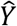 is an *N* × *K* matrix containing one vector of concentration and/or pH predictions per current response in *X′*_*test*_. We assess the performance of the regression by inspecting various qualities of the predictions such as:

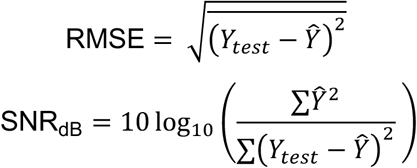

Additionally, we perform a multiple linear fit using the fitlm function in Matlab to evaluate the linearity of the predictions and check for any interactions among the neurotransmitter and/or pH predictions. We use these quality measures to compare RBS detection performance over different probes, different probe types, and different forcing functions.

### Resources

Source code (Matlab) and example *in vitro* datasets available upon request.

